# Detecting the anomaly zone in species trees and evidence for a misleading signal in higher-level skink phylogeny (Squamata: Scincidae)

**DOI:** 10.1101/012096

**Authors:** Charles W. Linkem, Vladimir Minin, Adam D. Leaché

## Abstract

The anomaly zone presents a major challenge to the accurate resolution of many parts of the Tree of Life. The anomaly zone is defined by the presence of a gene tree topology that is more probable than the true species tree. This discrepancy can result from consecutive rapid speciation events in the species tree. Similar to the problem of long-branch attraction, including more data (loci) will only reinforce the support for the incorrect species tree. Empirical phylogenetic studies often implement coalescent based species tree methods to avoid the anomaly zone, but to this point these studies have not had a method for providing any direct evidence that the species tree is actually in the anomaly zone. In this study, we use 16 species of lizards in the family Scincidae to investigate whether nodes that are difficult to resolve are located within the anomaly zone. We analyze new phylogenomic data (429 loci), using both concatenation and coalescent based species tree estimation, to locate conflicting topological signal. We then use the unifying principle of the anomaly zone, together with estimates of ancestral population sizes and species persistence times, to determine whether the observed phylogenetic conflict is a result of the anomaly zone. We identify at least three regions of the Scindidae phylogeny that provide demographic signatures consistent with the anomaly zone, and this new information helps reconcile the phylogenetic conflict in previously published studies on these lizards. The anomaly zone presents a real problem in phylogenetics, and our new framework for identifying anomalous relationships will help empiricists leverage their resources appropriately for overcoming this challenge.

---

The field of phylogenetics is poised to benefit tremendously from genomics, since resolving evolutionary relationships often requires massive amounts of data. Empirical phylogenetic researchers foresaw genomic data as a holy grail for resolving difficult relationships, such as rapid speciation events (Rokas et al. 2003; Dunn et al. 2008; Edwards 2009). However, phylogenetic conflict persists with genomic data, often with greater support, generating more questions than answers for many studies (Song et al. 2012; Gatesy and Springer 2014; Springer and Gatesy 2014; Pyron et al. 2014). In addition, reanalyses of previously published phylogenomic datasets often produce conflicting results (Dunn et al. 2008; Philippe et al. 2009, 2011), suggesting that analytical approach and model assumptions are critical to phylogenomic studies.

Processes such as horizontal gene transfer, gene duplication, and incomplete lineage sorting can lead to differences between species trees and gene trees (Maddison 1997). Species histories containing rapid diversification will have a high prevalence of incomplete lineage sorting due to few generations between speciation events. Rapid diversification in combination with a large effective population size can result in higher probabilities for gene trees that do not match the species tree than for gene tree that do match. These non-matching gene trees with high probability are referred to as anomalous gene trees (AGT), and the species tree branches that produce them are considered to be in the anomaly zone (Degnan and Rosenberg 2006). When the demographic history of the species tree is in the anomaly zone, sampling independent loci from the genome will result in AGT topologies being recovered at higher frequency than gene trees that match the species tree. Concatenation of these independent loci will result in strong support for the AGT topology whereas analyses using a species tree approach may recover the correct species tree (Kubatko and Degnan 2007; Liu and Edwards 2009).

Coalescent theory characterizes the anomaly zone for a four-taxon tree (Degnan and Rosenberg 2006) by showing that short internal branch lengths for an asymmetric topology will result in high probability for a symmetric AGT (Figure 1). The limit of the anomaly zone *a*(*x*) is defined by the following equation:

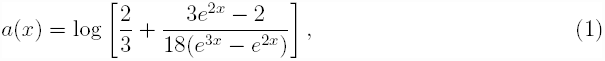

where *x* is the length of the branch in the species tree that has a descendant internal branch *y*. If the length of the descendant internal branch *y* is less than *a*(*x*), then the species tree is in the anomaly zone. As values of *x* get small, *a*(*x*) goes to infinity and therefore the value of the descendant branch *y* can be very long and still produce AGTs. In the four-taxon case the anomaly zone is limited by values of *x* greater than 0.27 coalescent units, when *a*(*x*) approaches zero, but this value increases with the number of taxa. In a four-taxon tree there is only one set of *x* and *y* internodes to consider and only three possible AGTs. With the addition of a single taxon, the five-taxon species tree has multiple sets of *x* and *y* and *z* internode branches that can have as many as 45 AGTs (Rosenberg and Tao 2008). The calculation of the multidimensional anomaly zone in trees larger than five-taxa is impractical, but a conservative simplification of the theory can be used for any species tree (Rosenberg 2013).

**Figure 1:**
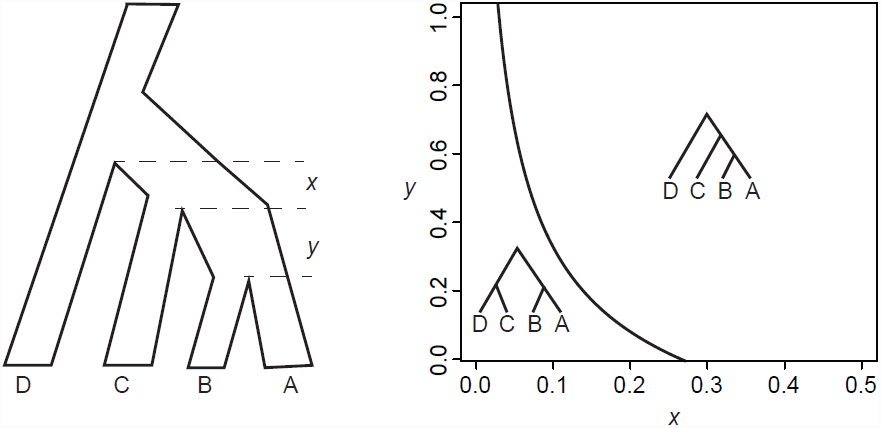
The length of branches X and Y in coalescent units in the species tree determine the probability of the gene tree topology. For branches under the anomaly zone curve the symmetric anomalous gene tree will have a higher probability than the asymmetric gene tree that matches the species tree.

Any species tree topology can be broken up into sets of four-taxon trees, which can individually be used in the anomaly zone calculation of equation (1). Rosenberg (2013) showed that focusing on sets of consecutive internal branches consistent with a four-taxon topology is a conservative estimate of the presence of the anomaly zone in any species tree. If the set of internodes fits the anomaly zone for the four-taxon case, at least one AGT exists, though more AGTs may occur due to nearby branches not considered in the isolated calculation (Rosenberg 2013). The unifying principle of the anomaly zone, that any four-taxon case within a larger phylogeny can be estimated independently, allows for the estimation of the anomaly zone in trees of any size.

The theoretical predictions of the anomaly zone are well characterized (Rosenberg and Tao 2008; Degnan and Rosenberg 2009; Rosenberg and Degnan 2010; Degnan et al. 2012a,b; Degnan 2013), and simulations have identified situations where certain phylogenetic methods succeed or fail under the anomaly zone (Steel and Rodrigo 2008; Huang and Knowles 2009; Liu and Edwards 2009; Liu et al. 2010b; Liu and Yu 2011). However, an empirical example of the anomaly zone has yet to be demonstrated. The lack of empirical evidence for the anomaly zone has led to doubt about the influence of the anomaly zone on real data (Huang and Knowles 2009), and the utility of coalescent methods for phylogenetic inference has been questioned (Gatesy and Springer 2014; Springer and Gatesy 2014). An investigation of the anomaly zone in an empirical setting requires an accurate species tree topology and estimates for ancestral branch lengths and population sizes, parameters that can be inferred accurately with hundreds of loci. Until recently, obtaining loci at this magnitude was not feasible for most non-model organisms, but new methods for obtaining large phylogenomic datasets are quickly changing the scale and scope of phylogenetic studies (Faircloth et al. 2012; McCormack et al. 2012; Song et al. 2012).

Here we present empirical evidence of the anomaly zone in a diverse radiation of lizards, the Scincidae. Using theoretical expectations, we define the set of species tree branches expected to generate anomalous gene trees based on the multispecies coalescent, and we apply these predictions to an empirical species tree. We use a new phylogenomic dataset collected using sequence capture of ultraconserved elements (Faircloth et al. 2012) and protein-coding genes (Wiens et al. 2012) to estimate the species tree, branch lengths, and population sizes required for identifying the anomaly zone in an empirical phylogeny.

## Scincidae

Scincidae is a large, diverse family of lizards found globally. The first division of higher-level skink relationships was largely based on skull morphology, dividing the family into four subfamilies: Acontinae, Felyninae, Lygosominae, and Scincinae (Greer 1970). Synapomorphies were defined for all but Scincinae, which was described as possessing the “primitive” form from which the other forms evolved, indicating Scincinae was not a natural group. Molecular phylogenies have largely supported the morphological hypotheses, although the relationships among the major groups within Scincinae have remained difficult to decipher. Multiple genetic studies have shown a pattern of relationships in which Acontinae is sister to all other Scincidae and that Felyninae and Lygosominae are nested within Scincinae (Whiting et al. 2003; Brandley et al. 2005, 2012; Wiens et al. 2012; Pyron et al. 2013; Lambert et al. 2014). Support for relationships within Scincinae and the placement of Lygosominae are low in all of these studies, despite sampling up to 44 genes (Wiens et al. 2012; Lambert et al. 2014) and up to 683 taxa (Pyron et al. 2013). These deep branches are often shown to be short when estimated from individual genes and concatenated genes, indicating diversification may have been rapid. Rapid diversification is an indication that specieation history may fit the demographic parameters consistent with the anomaly zone.

## MATERIALS AND METHODS

Identification of the causes of gene tree—species tree conflict in empirical phylogenies requires a large number of sampled genes on taxa that span difficult nodes (Liu and Edwards 2009). To accomplish this, we use a sequence capture next-generation sequencing approach to obtain 429 independent loci for all taxa of interest. The details of our data collection approach are below.

### Taxon sampling

Sampling was focused on fifteen species of skinks that span the deep nodes in the Scincidae tree where topological conflict is high. From the Scincinae subfamily we include *Brachymeles bonitae, Chalcides ocellatus, Eurylepis taeniolatus, Mesoscincus manguae, Ophiomorus raithmai, Plestiodon fasciatus,* and *Scincus scincus,* which represent the broad diversity in this difficult to resolve subfamily. These species have been used in prior studies of skink relationships based on Sanger sequencing genes (Brandley et al. 2005, 2012; Pyron et al. 2013) and have shown very short internode lengths and multiple alternative topologies suggesting anomolous gene trees may be present. A single Acontinae sample, *Typhlosaurus sp.,* is used to represent this well-supported subfamily (Lamb et al. 2010). We include at least one sample of four of the five groups in the Lygosominae (missing *Tiliqua): Mabuya unimarginata* for the *Mabuya* group; *Lygosoma brevicaudis* for the *Lygosoma* group; *Emoia caeruleocauda* for the *Eugongylus* group; and *Lobulia elegans, Sphenomorphus tridigitus, Sphenomorphus variegatus,* and *Tytthoscincus parvus* for the *Sphenomorphus* group. Previous studies (Honda et al. 2003; Reeder 2003; Skinner 2007; Skinner et al. 2011) have shown variation in the relationships between the *Lygosoma, Mabuya,* and *Eugongylus* groups, some with short internode lengths. A single outgroup taxon, *Xantusia vigils,* was chosen to root phylogenetic analyses.

### Probe design

We targeted a set of UCE loci combined with loci specific for squamates to maximize utility of our data in future studies of squamate phylogeny. We generated a set of sequencecapture probes based on a subset of the UCE probes from Faircloth et al. (2012) and some newly designed probes. We started with the *Tetrapods-UCE-5Kv1* probe set (www.ultraconserved.org) of 5,472 probes for Tetrapods and used blastn in CLC Genomics Workbench to screen probe sequences against a database of the *Anolis* genome (Alföldi et al. 2011) and a database of a whole-genome shotgun assembly of *Sceloporus occidentalis* (Harris and Leaché 2014). For this probe set, 1,125 probes matched the *Anolis* database and 1,554 probes matched the *S. occidentalis* database with 958 probes matching both databases. We then grouped the 958 probes by UCE and retained 2 probes for each locus. If a locus only had a single probe, we generated a new 120 bp probe with a 60 bp overlap to the existing probe. We excluded UCE loci that were within 100Kb of one another to reduce potential linkage. Consistent 2x tiling for all probes reduces potential capture bias, and can increase sequence capture efficiency over 1x probes (Tewhey et al. 2009).

To increase the relevance of our data for other squamate phylogeny studies, we developed probes for the 44 genes used in the squamate Tree of Life project (Wiens et al. 2012). Two 120 bp probes were designed for the center region of each gene, overlapping by 60 bp. In total, the probe set used for this study consists of 1,170 probes targeting 585 loci. Of those loci, 44 are commonly used in studies of squamates and 541 are UCE loci, common across all Tetrapods. This reduced set of sequence capture probes can be used across squamates. Probes were commercially synthesized into a custom MYbaits target enrichment kit (MYcroarray) and the probe sequences are available on Dryad (DOI).

### Library preparation, target enrichment, and sequencing

Whole genome extractions were completed for each sample using a *NaCl* extraction method (MacManes 2013). Genomic DNA (100 ng) was sonicated to a target peak of 400 bp (range 100–800 bp) using a Bioruptor Pico (Diagenode Inc.). Genomic libraries were prepared using an Illumina Truseq Nano library preparation kit with some minor modifications. Sonicated DNA was cleaned using a 2X volume of Agencourt AMPure XP beads and remained on the beads through the duration of the library preparation. Subsequent bead clean-up steps in the library preparation protocol consisted of adding an appropriate volume of 20% PEG-8000/2.5 M *NaCl* solution in place of adding more AMPure beads. Leaving the beads in solution reduces sample loss and cost of clean-up (Fisher et al. 2011). Final library bead clean-up used an 0.8X volume PEG solution to remove fragments smaller than 200 bp, which included an adapter-dimer produced during PCR enrichment.

Libraries were grouped into two sets of eight and pooled with equal concentration for 500 ng DNA per pool. Each pool was hybridized to the RNA-probes using the MYBaits kit with a modified protocol. We substituted a blocking mix of 500 uM (each) oligos composed of forward and reverse compliments of the Illumina Truseq Nano Adapters, with inosines in place of the indices, for the adapter blocking mix (Block #3) provided with the kit (oligo sequences on Dryad). We also substituted the kit supplied blocking mix #1 (Human Cot-1) for a chicken blocking mix (Chicken Hybloc, Applied Genetics Lab Inc.), which more closely matches our lizard targets. Library pools were incubated with the synthetic RNA probes for 24 hours at 65 °C. Post-hybridized libraries were enriched using Truseq adapter primers with Phusion taq polymerase (New England Biolabs Inc.) for 20 cycles. Enriched libraries were cleaned with AMPure XP beads. We quantified enriched libraries using qPCR (Applied Biosystems Inc.) with primers targeting five loci mapping to different chromosomes in the *Anolis* genome. The quality of enriched library pools was verified using an Agilent Tape-station 2200 (Agilent Tech.). These pools (and another pool of eight samples for another study) were pooled in equimolar ratio before sequencing. We sequenced all samples using a single lane of 150 bp, paired-end rapid-run sequencing on an Illumina HiSeq2500 at the QB3 facility at UC Berkeley.

### Preprocessing and de novo assembly

Data were processed using `Casava` (Illumina), which demultiplexes the sequencing run based on sequence tags. These raw data were then organized in a manner suitable for `Illumiprocessor v.2.0` (Faircloth 2013) using a series of scripts (github.org/cwlinkem/linkuce). Casava limits file sizes to 4 million reads for each individual, creating multiple files if sequencing coverage is extensive. These files were concatenated together into a single file for all forward reads and a single file for all reverse reads for each individual. Then all raw data files were transferred to a single folder to be processed by `Illumiprocessor`. A script (github) was used to create the configuration file based on a table of species names and the sequence file names. `Illumiprocessor` is a wrapper script to parallelize `Trimmomatic` (Bolger et al. 2014), which removes low-quality reads, trims low-quality ends, and removes adapter sequence. The cleaned paired-reads are organized by individual, based on the configuration file, and are ready for de novo assembly.

De novo assembly was conducted with the iterative de Bruijn graph short-read assembler, `IDBA` (Peng et al. 2010). This contiguous fragment (contig) assembler iterates over a set of k-mer values, removing the need to optimize the k-mer size as commonly done in other programs (eg. Velvet). We ran `IDBA` iteratively over k-mer values from 50 to 90 with a step length of 10.

### Dataset assembly

We used `phyluce` (Faircloth et al. 2012; Faircloth 2014) to assemble datasets of loci across taxa. We started by aligning species-specific contigs to the set of probes (match_contigs_to_probes.py) with `LASTZ` (Harris 2007). This script creates an `SQL` relational database of contig-to-probe matches for each taxon. We then query the database (get_match_counts.py) to generate a fasta file for the loci that are complete across all taxa. These data are aligned with `MAFFT` (Katoh and Standley 2013), and long ragged-ends are trimmed to reduce missing or incomplete data (seqcap_align_2.py). The final 429 loci that matched all taxa were exported in nexus format. Other file formats were obtained with `linkuce/file_converter.py`.

### Model testing, gene trees, and concatenation analyses

We used `modeltest_runner.py` to identify the models of substitution in the 95% confidence interval of the BIC for each locus using `jModelTest v2.1.5` (Guindon and Gascuel 2003; Darriba et al. 2012). The model with the lowest BIC score was chosen as the preferred model (Supplemental Information). Each locus was evaluated for the number of parsimony informative sites, number of constant sites, and number of variable sites using `PAUP* v.40b10` (Swofford 2003)(Supplemental Information). A maximum likelihood phylogenetic analysis was conducted on each locus using `RAxML v7.2.8` (Stamatakis 2006) with 1000 rapid-bootstrap replicates with the GTRGAMMA model.

All loci were concatenated (linkuce/concatenator.py) into a single alignment for maximum-likelihood (`RAxML`) and Bayesian analysis (`ExaBayes`) (http://sco.h-its.org/exelixis/web/software/exabayes/index.html). The concatenated data set was partitioned by locus (429 partitions) for both analyses. The ML analysis was conducted with the GTRGAMMA model with 1000 rapid-bootstrap replicates. `ExaBayes` analyses were run with the GTRGAMMA model with branch lengths linked across partitions and a parsimony starting tree with heated chains using different starting trees than the cold chain. Four independent runs were conducted, each with four chains, sampling every 500 generations. `ExaBayes` runs continued until the termination condition of mean topological difference less than 5% with at least 500,000 generations was met. Posterior distributions of trees were summarized with the consense script and posterior sample of parameters were assessed with `Tracer v1.5` (Rambaut and Drummond 2007) and combined with the `postProcParam script`.

### Species tree estimation

Due to the large number of genes sampled we limit our species tree estimation to a summary statistic approach. Species tree accuracy in summary statistic approaches is dependent on gene tree accuracy (Huang and Knowles 2009; Mirarab et al. 2014) since the methods rely solely on the structure of the fully resolved gene trees. Loci with few informative sites, often seen in NGS datasets, may not give strong support for all splits in the gene trees. This can potentially bias the species tree estimate and analyses relying on the species tree topology, such as the identification of the anomaly zone. We use the maximum pseudo-likelihood estimation of species trees `MP-EST v1.4` (Liu et al. 2010a) because it can accurately estimate the species tree topology despite the anomaly zone (Liu et al. 2010a). We accounted for gene tree and species tree uncertainty by running `MP-EST` using each iteration of the 1000 maximum-likelihood bootstrap replicates from `RAxML`. We created 1000 new tree files consisting of 429 trees, one bootstrap replicate of each locus. `MP-EST` was run on each of these files and an extended majority-rule consensus (eMRC) tree of the resulting species trees was calculated using `sumtrees in Dendropy` (Sukumaran and Holder 2010).

### Identifying anomalous nodes

We use the unifying principle of the anomaly zone (Rosenberg 2013) to determine which, if any, parts of the Scincidae eMRC species tree should produce AGTs. This procedure requires an estimate of internal branch lengths in coalescent units. We estimated branch lengths using two different methods. First, we used the branch lengths estimated by `MP-EST`, which jointly estimates internal branch lengths in coalescent units (based on *λ* = 2*τ*/*θ*) while maximizing the pseudo-likelihood of the species topology given the set of triplet topologies for each gene tree (Liu et al. 2010a, Equations: 6-8). Internal branches estimated by `MP-EST` may be shorter than expected when gene tree error is high (Mirarab et al. 2014), which may give an overestimation of internode pairs in the anomaly zone. Second, we estimated branch lengths using `BP&P v2. 1b` (Yang and Rannala 2010) with the original sequence data for 429 loci and the eMRC topology from `MP-EST. BP&P` uses a fixed species tree topology and the multi-species coalescent along with gene trees estimated using the Jukes-Cantor model to estimate branch lengths (*τ*) and population sizes (*θ*) (Rannala and Yang 2003; Burgess and Yang 2008). A gamma prior on *θ* (*α* = 2.0, *β* = 200) with a mean of 0.01 was used for populations size estimates on nodes. A gamma prior on *τ* (*α* = 4.0, *β* = 10.0) was used for the root node height with other times generated from the Dirichlet distribution (Yang and Rannala 2010). Rate variation between loci was accommodated with the random-rates model (Burgess and Yang 2008), in-which the average rate for all loci is fixed at 1 and the rates among loci are generated from a Dirichlet distribution. We used an *α* of 2.0 for moderate variation among loci. The MCMC chain was run for 100,000 samples, sampling every 10 generations for a total of 1,000,000 sampled states with a burnin of 150,000 states. Three independent analyses were conducted to verify convergence on a stable posterior. `BP&P` results were converted to coalescent units (*λ* = 2*τ*/*θ*) consistent with those calculated by `MP-EST`.

Each pair of parent-child internodes were compared to the anomaly zone based on values of *λ* calculated from `BP&P` and `MP-EST`. The value for the parent nodes (*x*-nodes) were put into equation (1) for the limit of the anomaly zone in a four-taxon asymmetric tree to determine if they are inside the zone and would therefore produce anomalous gene trees. If the value of the child (*y*-node) is less than *a*(*x*), the pair of internodes are in the anomaly zone and AGTs are expected. This calculation was first conducted on the median values of branch lengths from the eMRC tree of the `MP-EST` species tree replicates and the median values of τ and *θ* from the eMRC tree of the `BP&P` posterior distribution. Additionally, for the `MP-EST` bootstrap replicates, anomaly zone calculations were done for each internode pair for each species tree bootstrap replicate, accounting for topological error in estimates of branch lengths. For `BP&P`, 1000 random draws of joint values of *τ* and *θ* for the internode pairs were made from the posterior distribution and compared to the anomaly zone. The `BP&P` analysis is only performed on the eMRC topology. Scripts to perform these function relied on the Dendropy package (Sukumaran and Holder 2010). We report the proportion of bootstrap replicates that match the eMRC tree and are in the anomaly zone with `MP-EST` and the proportion of the 1000 draws from `BP&P` that are in the anomaly zone.

## RESULTS

### Genomic data and assemblies

All samples were successfully sequenced at sufficient levels to result in high coverage of target loci. Some samples represented a larger portion of the sequencing (Table 1) potentially due to unequal pooling prior to hybridization. Raw reads averaged 9.8 million (range 5.4–21.6) reads per species with most reads being high-quality, resulting in a low rate of trimming and removal. Sequencing resulted in higher coverage than needed for the sequence capture approach due to using 24 samples on the sequencing lane instead of the full potential of 96 samples. The added sequence coverage resulted in a higher proportion of off-target sequencing than would be expected, including complete mitochondrial genomes for many taxa. Results from other experiments have shown that off-target sequencing is reduced when more samples are included in the sequencing lane without a loss of target sequence coverage (unpublished data).

**Table 1:** Genomic data collected and assembly results for 15 species of skinks and the out group.

| Species | Voucher | Total reads<br>(Million) | Clean reads<br>(Million) | IDBA<sup>a</sup> |  |
| --- | --- | --- | --- | --- | --- |
|  |  |  |  | Contigs | Loci |
| *Brachymeles bonitae* | AJB077<sup>c</sup> | 8.71 | 7.71 | 29,942 | 550 |
| *Chalcides ocellatus* | MVZ242790<sup>b</sup> | 6.59 | 5.93 | 26,745 | 569 |
| *Emoia caeruleocauda* | KU307155<sup>c</sup> | 6.08 | 5.64 | 21,414 | 563 |
| *Eurylepis taeniolatus* | MVZ246017<sup>b</sup> | 7.42 | 6.72 | 30,163 | 572 |
| *Lobulia elegans* | BPBM18690<sup>d</sup> | 10.2 | 9.44 | 49,859 | 556 |
| *Lygosoma brevicaudis* | MVZ249721<sup>b</sup> | 8.40 | 7.70 | 47,774 | 553 |
| *Mabuya unimarginata* | CWL615<sup>e</sup> | 19.5 | 17.9 | 134,666 | 533 |
| *Mesoscincus manguae* | CWL614<sup>e</sup> | 11.4 | 10.5 | 38,828 | 550 |
| *Ophiomorus raithmai* | MVZ248453<sup>b</sup> | 7.78 | 7.00 | 35,506 | 568 |
| *Plestiodon fasciatus* | KU289464<sup>c</sup> | 12.6 | 11.7 | 67,908 | 542 |
| *Scincus scincus* | MVZ234538<sup>b</sup> | 6.54 | 5.94 | 20,231 | 566 |
| *Sphenomorphus tridigitus* | FMNH258830<sup>f</sup> | 8.39 | 7.82 | 39,350 | 560 |
| *Sphenomorphus variegatus* | KU315087<sup>c</sup> | 8.99 | 8.31 | 43,927 | 550 |
| *Typhlosaurus sp* | MVZ164850<sup>b</sup> | 8.35 | 7.60 | 31,261 | 549 |
| *Tytthoscincus parvus* | JAM6275<sup>b</sup> | 5.41 | 5.00 | 18,850 | 568 |
| *Xantusia vigilis* | KU220092<sup>c</sup> | 21.5 | 19.9 | 155,978 | 550 |

<sup>a</sup> Iterative de Bruijn graph short-read assembler.
<sup>b</sup> Museum of Vertebrate Zoology, UC Berkeley, CA.
<sup>c</sup> University of Kansas, Lawrence KS.
<sup>d</sup> Bernice Pauahi Bishop Museum, Honolulu, HI.
<sup>e</sup> No voucher specimen. Tissue deposited at KU.
<sup>f</sup> Field Museum of Natural History, Chicago IL.

The number of contigs found for `IDBA` for each individual is large, averaging over 49,000 (range 18,850–155,978) contigs across the 16 species. The `IDBA` assemblies match most of the 585 loci targeted. Datasets were assembled for complete taxon sampling for all loci. `IDBA` assemblies resulted in 429 loci across all taxa. Of the 44 loci used in previous squamate systematics studies, only two loci were present in `IDBA` assemblies for all species.

### Loci informativeness and model choice

The 429 loci from the `IDBA` assembly totals 276,480 nucleotide positions with 5.28 % missing data and an average length of 644 base-pairs (range 338–1070). Individual loci vary in character variability with an average of 6% parsimony informative sites (range 0–15%) and 18% variable sites (range 1–48%). Most loci have a best-fit model matching HKY or K80 (400 out of 429) suggesting a prevalence of transition/transversion bias in these genomic loci (Table 2). Model testing shows a preference for either a gamma or invariant-sites model (376 out 429) but the combination is rarely preferred (38 out of 429). Only 18 loci have a preferred model that does not accommodate rate-heterogeneity (Table 2).

**Table 2:** Summary of model testing results ranked by model complexity. The model chosen for each locus can be found in the supplemental information.

| Model | # free parameters | # of Loci | Avg % informative | Avg % variable |
| --- | --- | --- | --- | --- |
| K80 | 31 | 1 | 0.8 | 4.5 |
| K80+I | 32 | 13 | 5.1 | 14.5 |
| K80+G | 32 | 22 | 6.9 | 21.3 |
| K80+I+G | 33 | 9 | 8.0 | 22.3 |
| F81 | 33 | 3 | 0.2 | 1.6 |
| F81+G | 34 | 3 | 2.3 | 8.6 |
| HKY | 34 | 11 | 1.2 | 7.8 |
| HKY+I | 35 | 111 | 3.9 | 13.2 |
| HKY+G | 35 | 209 | 6.3 | 21.1 |
| HKY+I+G | 36 | 26 | 6.5 | 20.0 |
| SYM+I | 36 | 1 | 6.6 | 19.1 |
| SYM+G | 36 | 3 | 8.6 | 23.7 |
| GTR+G | 39 | 13 | 7.5 | 22.0 |
| GTR+I | 39 | 1 | 6.1 | 16.5 |
| GTR+I+G | 40 | 3 | 8.4 | 23.6 |

### Gene trees

Maximum Likelihood searches for individual loci resulted in 429 unique topologies, one for each locus. These topologies also differ from the concatenation ML tree and the species tree. Bootstrap replicates average 925 (range 163–1000) unique topologies out of the 1000 replicates indicating that gene tree resolution is low for most individual loci.

Concatenated gene trees are largely congruent between the ML and Bayesian runs (Figure 2). Acontinae is sister to all other taxa with strong support (100 bootstrap and 100 posterior probability). Lygosominae is monophyletic and the *Sphenomorphus* group is sister to the clade of the *Mabuya, Lygosoma,* and *Eugongylus* groups. The *Eugongylus* group is sister to the *Lygosoma* group. All of these relationships have strong support in both analyses. There is also strong support for *Brachymeles* to be sister to Lygosominae, making Scincinae paraphyletic. The only topological difference between the two analyses is the placement of *Ophiomorus*. In the ML analysis *Ophiomorus* is sister to the other genera in Scincinae (minus *Brachymeles)* with a bootstrap score of 51, whereas in the Bayesian analysis *Ophiomorus* is sister to all Scincinae and Lygosominae with a posterior probability of 71. In both cases the support for the placement of *Ophiomorus* is low despite the large amount of data used in these analyses.

**Figure 2:**
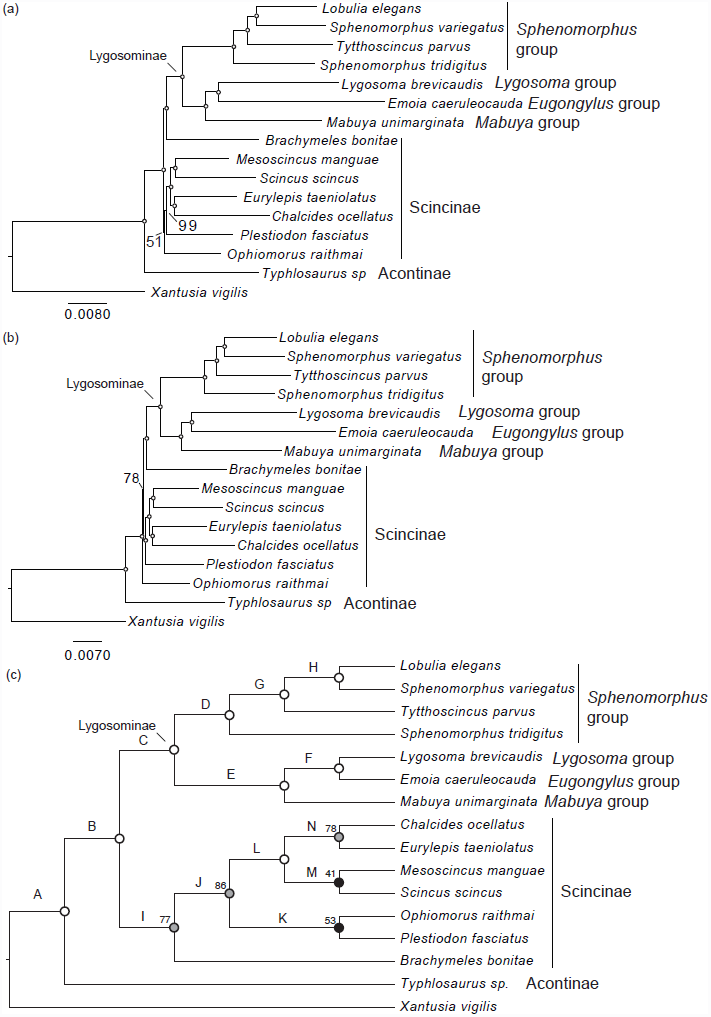
Majority-rule consensus trees from the concatenated loci run in `RAxML` (a), `ExaBayes`, (b) and the species tree from `MP-EST` (c). Open circles indicate 100% bootstrap support or a Bayesian posterior probability of 1.0. Nodes with lower support are labelled. Both concatenation analyses have similar topologies with differences in the placement of *Ophiomorus* and the `RAxML` tree is slightly longer. The species tree (c) is shown as a cladogram. Relationships within the Lygosominae are the same across analyses. The relationships within Scincinae differ both between the concatenation analyses and in comparison to the species tree. Letters on species tree nodes are used for Table 3 and discussion of internode pairs.

### Species trees

There were 110 unique species tree topologies found using the bootstrap replicates. The `MP-EST` eMRC species tree has 100% support for many relationships (Figure 2c), but some key relationships are poorly supported and differ from the concatenation tree. Most significantly, the placement of *Brachymeles* in the species tree recovers a monophyletic Scincinae, but with low (bootstrap = 77) support. The sister relationships of *Scincus* and *Mesoscincus* has the lowest support (bootstrap = 48). *Ophiomorus* is sister to *Plestiodon* with low support (bootstrap = 53). The Lygosominae portion of the species tree is identical to the concatenation topologies.

### Nodes in the anomaly zone

*MP-EST*. —Median internode lengths calculated with `sumtrees` (Sukumaran and Holder 2010) were used to calculate the anomaly zone for each pair of internodes in the eMRC (equation (1)). A large region of the phylogeny is in the anomaly zone based on these median values (Figure 3a). The majority of relationships in the Scincinae subfamily have internode lengths that are expected to produce AGTs. Examining the anomaly zone across the species tree replicates (Figure 3b) shows that most pairs of parent-child internodes found to be in the anomaly zone with median branch lengths remain in the anomaly zone across replicates. One region of the phylogeny leading to the Lygosominae (yellow branches) is only in the anomaly zone for 6% of the replicates that match the eMRC topology. The species tree replicates include relationships not found in the eMRC that represent the alternate resolutions of the poorly resolved nodes. Many of these alternative species tree relationships in the replicate set of trees are also in the anomaly zone (results not shown).

**Figure 3:**
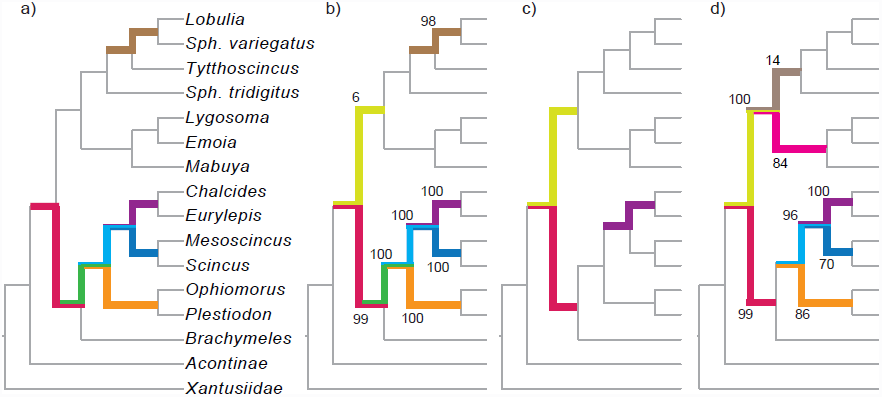
Majority-rule consensus topology shown as a cladogram. Pairs of internodes that are under the anomaly zone curve are highlighted in bold colors on each tree. Tree a) is based on median values of branch lengths calculated from MP-EST. Tree b) shows the the frequency of the internodes in the anomaly zone across bootstrap replicates that match the eMRC topology. Tree c) shows the occurrence based on the median values of branch lengths from the posterior distribution of BP&P. Tree d) shows the frequency of the internode in the anomaly zone based on 1000 draws of joint internode values from the posterior distribution.

*BP&P*.—The posterior distribution from `BP&P` was summarized using sumtrees to obtain the median branch length *τ* and population size *θ* for internodes. `BP&P` calculates *τ* and *θ* individually, which were used to calculate *λ* (*λ* = 2*τ*/*θ*) (Table 3). Internodes B, C, and N (Figure 2c) are particularly short and internode G is the longest. Calculations of *a*(*x*) (equation (1)) show that six internodes (B, C, J, K, L, N) have *λ* values above zero indicating they may produce AGTs depending on the length of the descendant internode. Of these, B, C, J, and L have descendant internodes. The internode pairs B/C, B/I, and L/N are in the anomaly zone because the lengths of internodes C and I are shorter than *a*(*x*) for B and internode N is shorter than *a*(*x*) for L (Figure 3c). To account for the range in branch length estimates across the posterior distribution, 1000 random draws of *τ* and *θ* were made. Each draw was calculated for occurrence of the anomaly zone for each pair of internodes. Internodes inferred to be in the anomaly zone using median branch lengths were all inferred at high frequency (Figure 3d). Additional pairs of internodes are in the anomaly zone when taking into account the range of *τ* and *θ* values in the posterior distribution, showing a similar pattern of anomaly zone nodes as found with estimates from `MP-EST`.

**Table 3:** Median branch lengths (*τ*) and population sizes (*θ*) used to calculate the coalescent unit (*λ*) for the nodes labelled on the `MP-EST` species tree (Figure 2c). *a*(*x*) indicates the limit of the anomaly zone for that node length. If the length of the descendant node is smaller than *a*(*x*), then the node pair is in the anomaly zone. If *a*(*x*) is < 0 the node pair will not produces anomalous trees. Descendant nodes in bold indicate anomaly zone pair.

| Node | $\tau$ | $\theta$ | $\lambda$ | $a(x)$ | Descendant node(s) |
| --- | --- | --- | --- | --- | --- |
| A | 0.006391 | 0.016352 | 0.781678 | $< 0$ | B |
| B | 0.000036 | 0.012459 | 0.005778 | 1.020 | **C, I** |
| C | 0.000117 | 0.008112 | 0.028846 | 0.443 | D, E |
| D | 0.001574 | 0.00686 | 0.45889 | $< 0$ | G |
| E | 0.002552 | 0.007271 | 0.701966 | $< 0$ | F |
| F | 0.001553 | 0.01005 | 0.309054 | $< 0$ | — |
| G | 0.007702 | 0.007601 | 2.02657 | $< 0$ | H |
| H | 0.001286 | 0.00518 | 0.49652 | $< 0$ | — |
| I | 0.005576 | 0.014087 | 0.79165 | $< 0$ | J |
| J | 0.001754 | 0.024723 | 0.14189 | 0.0832 | L, K |
| K | 0.000902 | 0.013989 | 0.128958 | 0.0982 | — |
| L | 0.000588 | 0.010911 | 0.10778 | 0.1284 | M, **N** |
| M | 0.001272 | 0.00959 | 0.26527 | 0 | — |
| N | 0.000154 | 0.012777 | 0.024105 | 0.4994 | — |

## DISCUSSION

Species trees often differ from concatenated gene trees, and the anomaly zone may be the culprit causing these disagreements. The anomaly zone could also explain why some phylogenomic studies find low support for relationships despite the inclusion of hundreds to thousands of loci. In the empirical example of skinks presented here, we find strong conflict between species trees and concatenated gene trees, as well as conflict between individual gene trees. This type of conflict is typical for phylogenomic studies of rapid diversification events (Zou et al. 2008; McCormack et al. 2012). Our examination of the anomaly zone in skinks shows that the parts of the tree in conflict correspond with areas of the tree that are also in the anomaly zone. The anomaly zone is a potential explanation for why conflicting relationships persist in some phylogenomic studies.

### The anomaly zone in empirical phylogenies

We find that the anomaly zone likely occurs in empirical studies and that it may be more pervasive than previously assumed (but see Huang and Knowles (2009)). Although there might be insufficient variation to resolve gene trees that are derived from a species tree in the anomaly zone, most studies are not specifically interested in whether or not we can estimate individual gene trees. The species tree is the target of analysis, not the individual gene trees (Edwards 2009). While it is true that estimation of any particular gene tree is hindered by low genetic variation, this does not change whether or not speciation events occurred quickly enough to place them into the anomaly zone. Gene tree analyses of concatenated loci, as a proxy for the species tree, can result in strong support as more data are added, even if there is low genetic variation in individual loci. If the speciation history is in the anomaly zone, then the resulting phylogeny will be erroneous (Kubatko and Degnan 2007). Our study shows that there is low support for relationships in most individual gene trees, but after concatenating the 276,480 characters they provide strong support for most relationships (Figure 2a–b). Huang and Knowles (2009) showed that when the species history is in the anomaly zone, gene tree estimation error will be high and that the lack of variation will make estimation of individual gene trees difficult. The observation of significant gene tree discordance across hundreds of loci may be a good sign that the species history is the anomaly zone, and the framework that we provide here offers one way to test this hypothesis.

Summary method for species tree inference have become a necessity, since the more statistically rigorous full-Bayesian approaches (*BEAST and BEST) cannot handle hundreds of loci (Bayzid and Warnow 2013). These summary methods use gene trees to estimate species trees, and as previously discussed, gene tree estimation error is typically high in cases where an anomaly zone is suspected. Gene tree estimation error reduces the accuracy of species tree estimation (Bayzid and Warnow 2013; Mirarab et al. 2014), which makes species tree inference in the anomaly zone more difficult (Liu and Edwards 2009).

We accounted for gene tree error by repeating the species tree estimation procedure using the gene trees constructed from the bootstrap replicates, which can provide a measure of accuracy that is not available when only using the ML gene trees (Mirarab et al. 2014). Despite this, it is possible that due to gene tree error our species tree estimate is not correct, but any inaccuracies should be reflected by the low bootstrap support for the species tree. By estimating the anomaly zone across all topologies in the bootstrapped species tree from `MP-EST` we show that the anomaly zone can be inferred even when species trees are not entirely resolved. Species trees in the anomaly zone are likely to have nodes with low bootstrap support due to the frequency of AGTs and the occurrence of gene tree error. In these situations, increased gene sampling may not increase node support due to the addition of more gene tree estimation error. Increasing the accuracy of gene trees through sampling longer loci (McCormack et al. 2009) or by combining loci with a shared history into larger gene fragments (Bayzid and Warnow 2013; Betancur-R et al. 2013) may improve inference of species trees in the anomaly zone.

### Frequency of anomalous gene trees

While we cannot be certain that the skink species tree that we estimated is correct, it is clear that a set of species tree topologies estimated from the phylogenomic data shows signs of the presence of the anomaly zone over multiple pairs of internodes (Figure 3). The extent and frequency of the anomaly zone in this empirical example indicates that many AGT topologies may exist across the genomes of these taxa. It is important to keep in mind that each inference of the anomaly zone is limited to pairs of internal branches, without consideration of neighboring relationships. This simplification can be made because accounting for other branches can only increase the size of the anomaly zone (Rosenberg and Tao 2008), making our inference a conservative approximation of the extent of AGTs. When considered together, the extent of the anomaly zone in our empirical example has the potential to produce many AGTs.

The expectation in the four-taxon case is a symmetric AGT, with as many as three AGTs when both branches are very short (Figure 1). With a single occurrence of the anomaly zone in a tree of four taxa it would be easy to predict the shape of the AGT and therefore the expected shape of the concatenation tree. With more taxa the ability to predict the anomalous result disappears due to the anomaly zones spanning multiple sets of nodes or when other short branches are near the anomaly zone. Rosenberg and Tao (2008) showed that the number of AGTs increases rapidly as the number of short internodes increases and that an anomaly zone that includes three internodes (five-taxon trees or larger) can produce as many as 45 AGTs. Estimates for trees larger than five taxa or an anomaly zone spanning more than three nodes have not been estimated, but are expected to increase exponentially. Simulation studies testing the ability of different species tree methods to overcome the anomaly zone largely focus on small trees with a single pair of internodes in the anomaly zone (Kubatko and Degnan 2007; Huang and Knowles 2009; Liu and Edwards 2009). It is unclear how species tree methods will perform under multiple sets of anomalous internodes in either close proximity to one another, or spread throughout the tree as we see in skinks. Additionally, with a large anomaly zone a “Wicked forest” may occur, in which the AGT will be the same topology as an alternative species tree topology that also produces AGT of other topologies (Degnan and Rosenberg 2006; Rosenberg and Tao 2008). The possibility that species tree methods can estimate the correct species tree in cases of a wicked forest are unknown.

Edwards (2009) predicted that phylogenomic studies would find lower support for relationships using species tree methods than would be obtained from concatenation, especially in older clades. This prediction is based on the ideas that missing data would have a larger effect on species trees, and that species trees use more complex models compared to concatenation. We find these predictions to be accurate, though their cause may be different than originally proposed. We propose that low support in species tree analyses is due to a combination of gene tree estimation error and inherent properties of the speciation process during rapid diversification producing multiple AGT topologies that bias phylogenetic signal. Simulation studies have shown that many species tree methods can overcome the anomaly zone in simple four-taxon scenarios (Liu and Edwards 2009), but no study has looked at the effect of larger anomaly zone problems on trees with more taxa. We predict that when the anomaly zone occurs across more than two internodes, the greater number of AGTs will provide support for multiple species trees, reducing the support for some species tree nodes. This will likely occur even when gene trees are estimated with certainty (i.e., using simulated data), a luxury not available with empirical studies. The increased number of AGTs may also result in low support in concatenation analyses even when analyzing hundreds of genes if the AGTs are in direct topological conflict. In this study, we find two nodes with `RAxML` and a different node in `ExaBayes` that have low support despite having over 276,000 characters in the analyses. With multiple AGTs there may be alternative topologies with high probability in the set of candidate genes. These alternative topologies should lower branch support in concatenation analyses. This low support for nodes near short branches may be an indication of an anomaly zone problem in phylogenomic datasets.

### Higher level skink relationships

Resolving the relationships within Scincidae is an ongoing challenge (Brandley et al. 2012; Wiens et al. 2012; Pyron et al. 2013; Lambert et al. 2014) which our current study addresses with a slightly different approach. Previous studies have used hundreds of taxa (Pyron et al. 2013) or many loci (Wiens et al. 2012; Lambert et al. 2014) to try and resolve the relationships in this large, diverse family of lizards. Our taxon sampling is most similar to Brandley et al. (2012) but utilizes 429 loci and species tree analyses to estimate relationships. Our preferred estimate of species relationships (Figure 2c) shows subfamily relationships concordant with results in Pyron et al. (2013) and Lambert et al. (2014). Acontinae is sister to all other skinks and the subfamilies Scincinae and Lygosominae are monophyletic. The monophyly of Scincinae conflicts with most studies and our own results from analyses of concatenated data (Whiting et al. 2003; Brandley et al. 2005, 2012; Wiens et al. 2012) but is likely the more accurate relationship based on our inference of the anomaly zone in relation to the nodes preceding and in this subfamily. Similar to Lambert et al. (2014) we find *Brachymeles* to be the sister genus to all other Scincinae as opposed to sister to all Lygosominae (Brandley et al. 2012). The relationships among Scincinae genera differs from those presented in Brandley et al. (2012) though we have far fewer genera sampled and low support so many comparisons should be made with caution. A detailed examination of the relationships among the genera in the Scincinae with broad taxon sampling is clearly warranted. Our inference of the anomaly zone among some of the Scincinae genera suggests that hundreds of loci and species tree analyses will be necessary to accurately estimate the phylogenetic relationships within this group.

Relationships within the Lygosominae are largely concordant with previous studies in finding the *Sphenomorphus* group sister to all other groups (Honda et al. 2003; Reeder 2003; Skinner et al. 2011). We find the *Lygosoma* group to be sister to the *Eugongylus* group and that this pair is sister to the *Mabuya* group consistent with the results of Reeder (2003) and Skinner et al. (2011). Relationships among the sampled genera in the *Sphenomorphus* group are consistent with previous results (Linkem et al. 2011). Over three-quarters of all skink species are in the Lygosominae and it appears based on our limited sampling that the broad groupings of genera can be consistently resolved and the anomaly zone is not an issue at this level. Within the *Sphenomorphus* group, previous studies have reported short branches separating major groups (Linkem et al. 2011; Skinner et al. 2011). We suspect that there will be anomaly zone issues within the *Sphenomorphus* radiation. The *Eugongylus* group will likely also present anomaly zone issues given the large number of species in the group and relatively recent origin. Further work on these diverse groups is needed to better understand their systematic relationships.

### Overcoming the anomaly zone

Sequencing hundreds of unlinked loci provides an opportunity to explore the conflicts between loci and analytical approaches, as well as address what may be the source of conflict. Our work shows that researchers conducting empirical studies should closely consider the potential impact the anomaly zone has on their phylogenetic analyses. A common trend with phylogenomic studies is to analyze the data with concatenation, an approach that has the advantage of faster computation times and simplicity, but that provides overwhelmingly and likely erroneous strong support across most the tree. Species tree analyses often result in lower support for difficult parts of the tree than concatenation. Instead of marginalizing the species tree results, we should acknowledge that they are a likely consequence of the speciation history for the group. The lower support provided by coalescent-based species tree inference is potentially a more accurate reflection of the support for the tree given the data.

As we show here, the anomaly zone is likely more pervasive than previously suggested and should be accounted for when studying taxa that may have diverged rapidly, even if that rapid event was in the distant past. Combining hundreds to thousands of independent loci together with coalescent-based species tree inference is the most effective way of getting an accurate result. Targeting longer loci will help reduce gene tree estimation error, resulting in a better estimate of the species tree. In the most extreme cases, resolving the nodes of a species tree with strong support may not be possible even when sampling the entire genome.

## ACKNOWLEDGEMENTS

We would like to thank R. Brown of the University of Kansas, C. Spencer and J. McGuire of UC Berkeley, A. Resetar of the Field Museum, and K. Imada and A. Allison of the Bishop Museum for loans of tissues necessary for this work. B. Faircloth provided detailed help in troubleshooting early attempts at sequence capture, and we thank him for quickly responding to our queries. R. Harris helped with designing this new probe set. R. Bryson provided useful feedback and discussion. C.W.L. was supported by an NSF Postdoctoral Research Fellowship in Biology (Award 1202754). Data collection was funded by NSF grant DBI-1144630 awarded to A.D.L. This work used the Vincent J. Coates Genomics Sequencing Laboratory at UC Berkeley, supported by NIH S10 Instrumentation Grants S10RR029668 and S10RR027303.

